## Supplementary figures and images for "Genetic analysis of F1 cluster phages that infect *Mycobacterium smegmatis* identifies two distinct holin-like proteins that regulate the host lysis event"

### S2 Figure 1

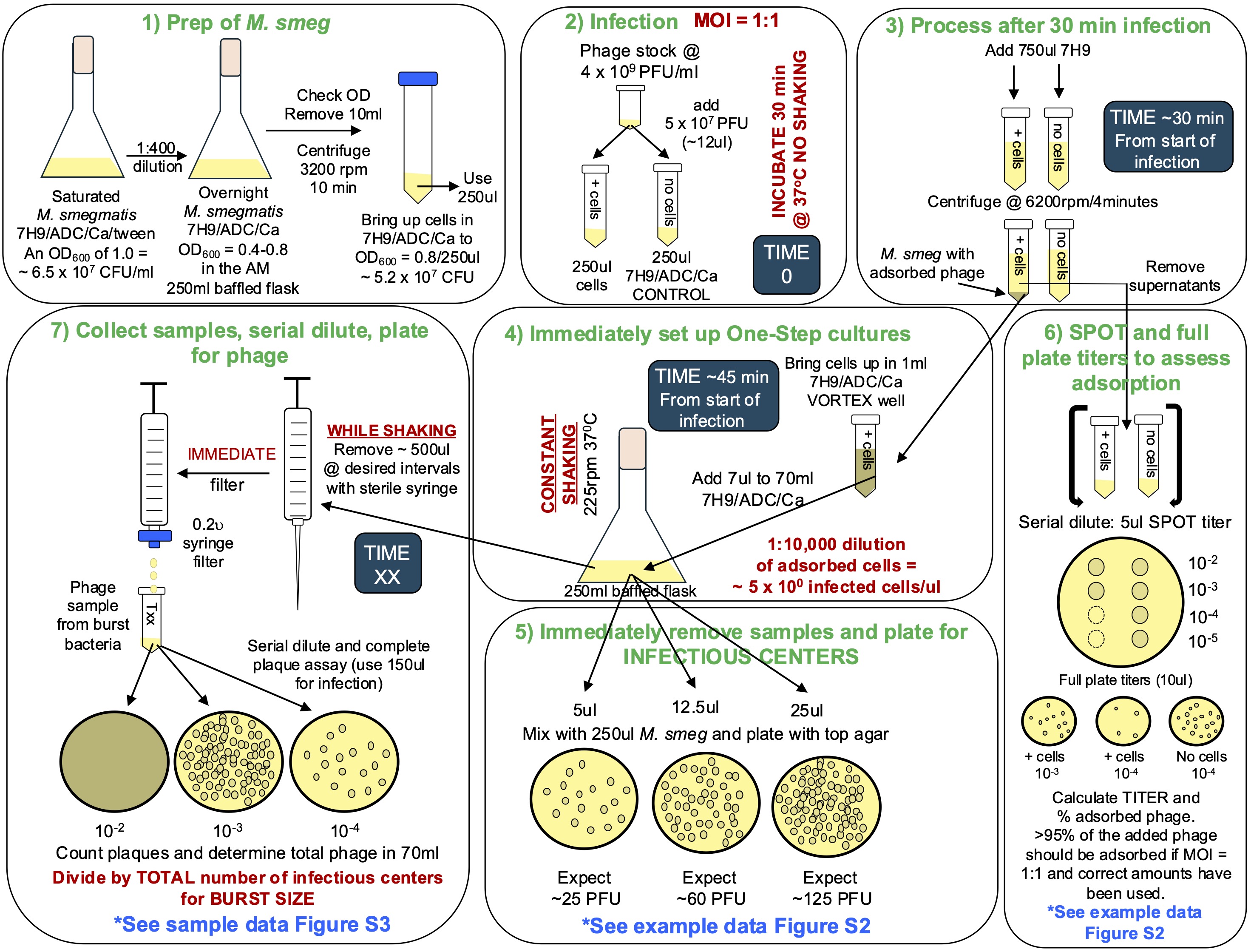

### S3 Figure 2

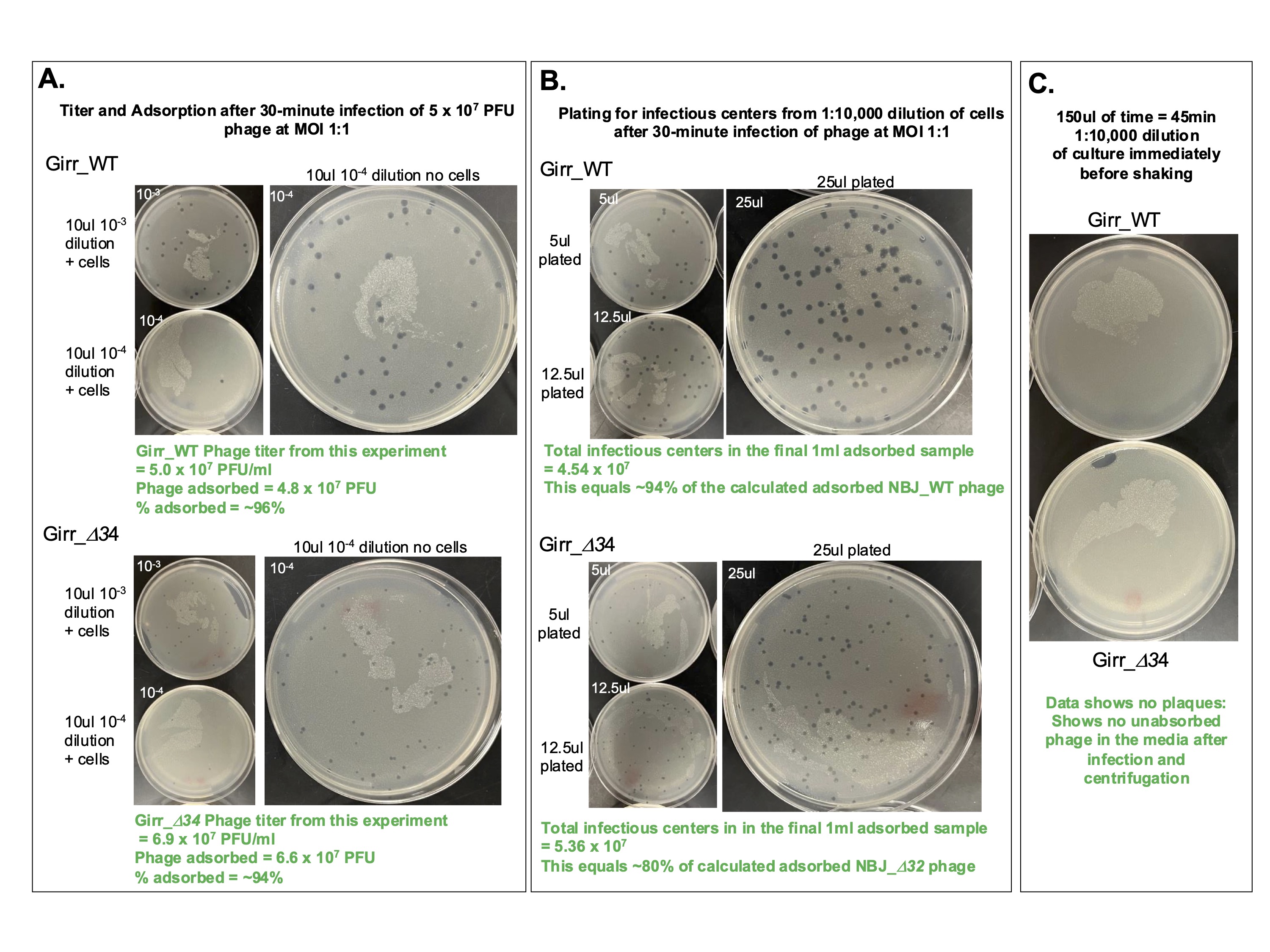

### S4 Figure 3

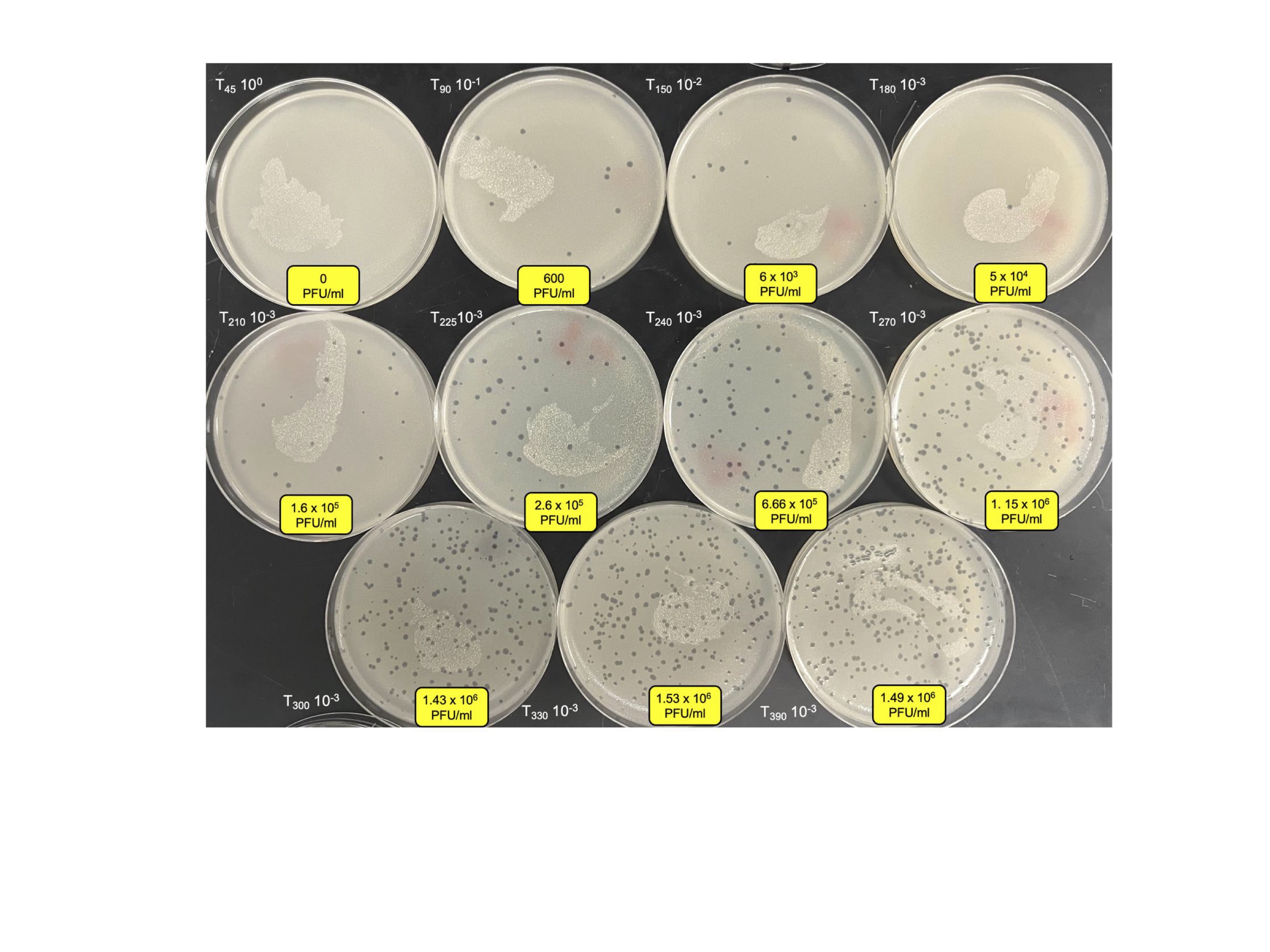

### S5 Figure 4

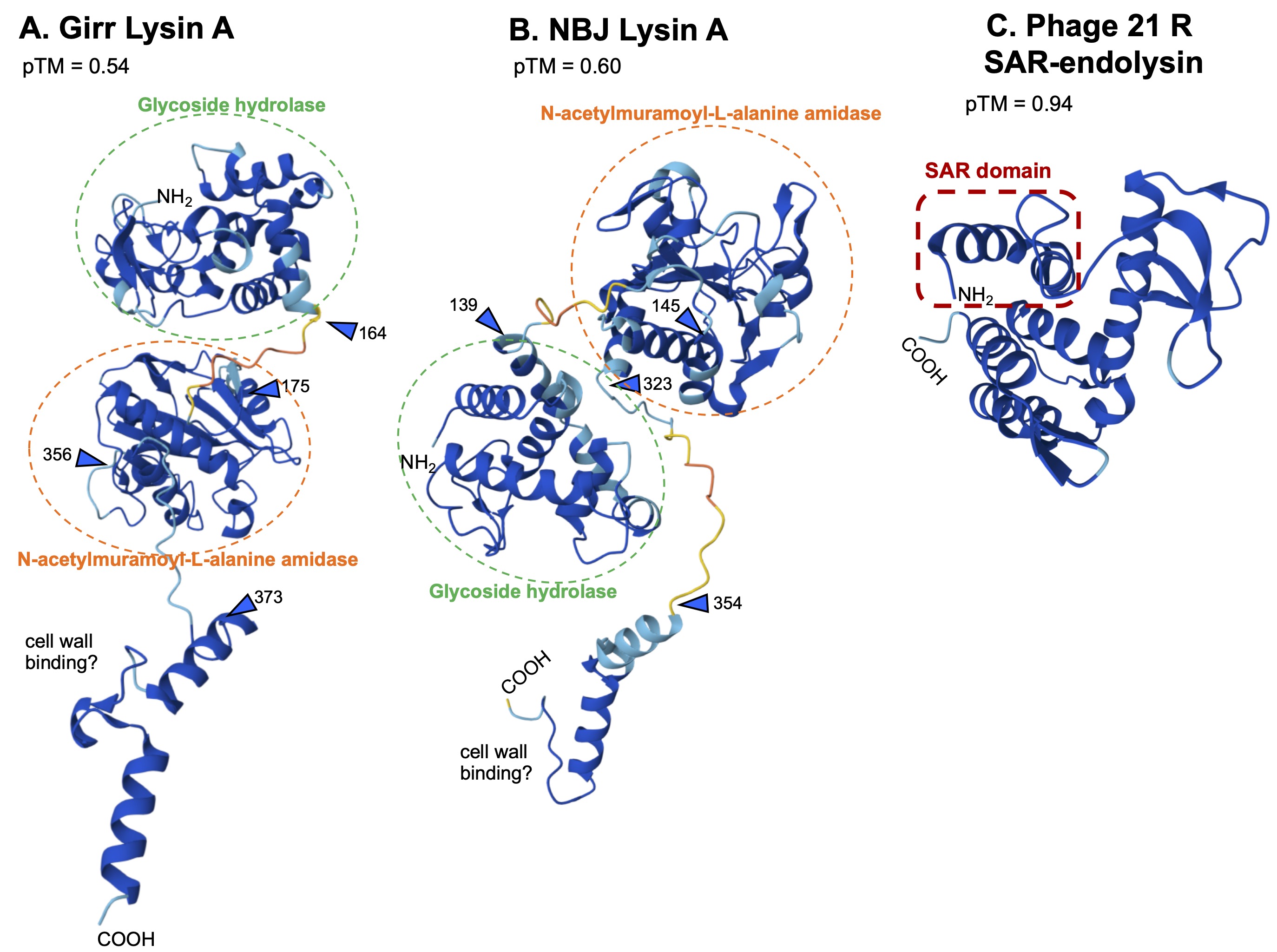

### S6 Figure 5

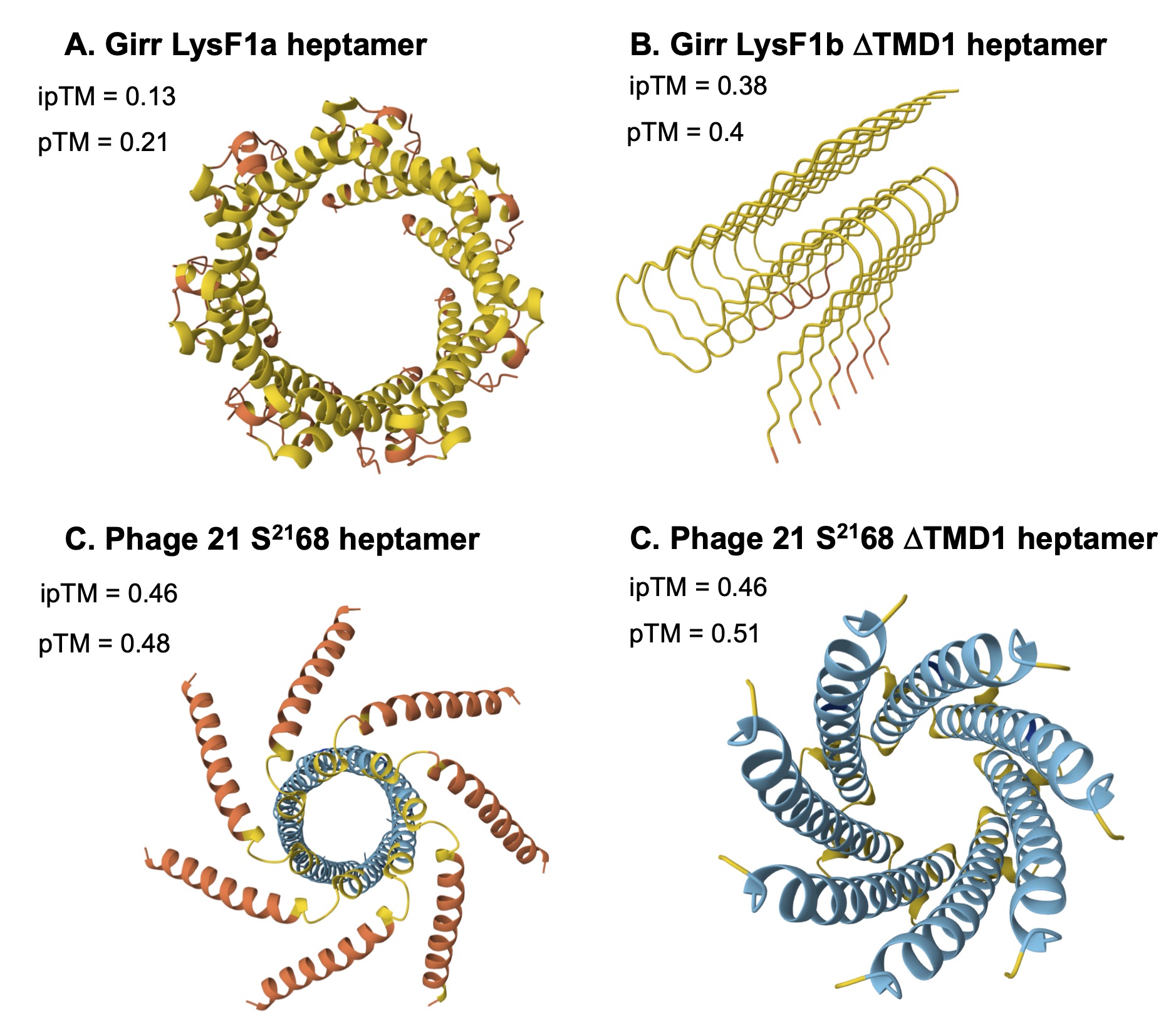

### S7 Figure 6

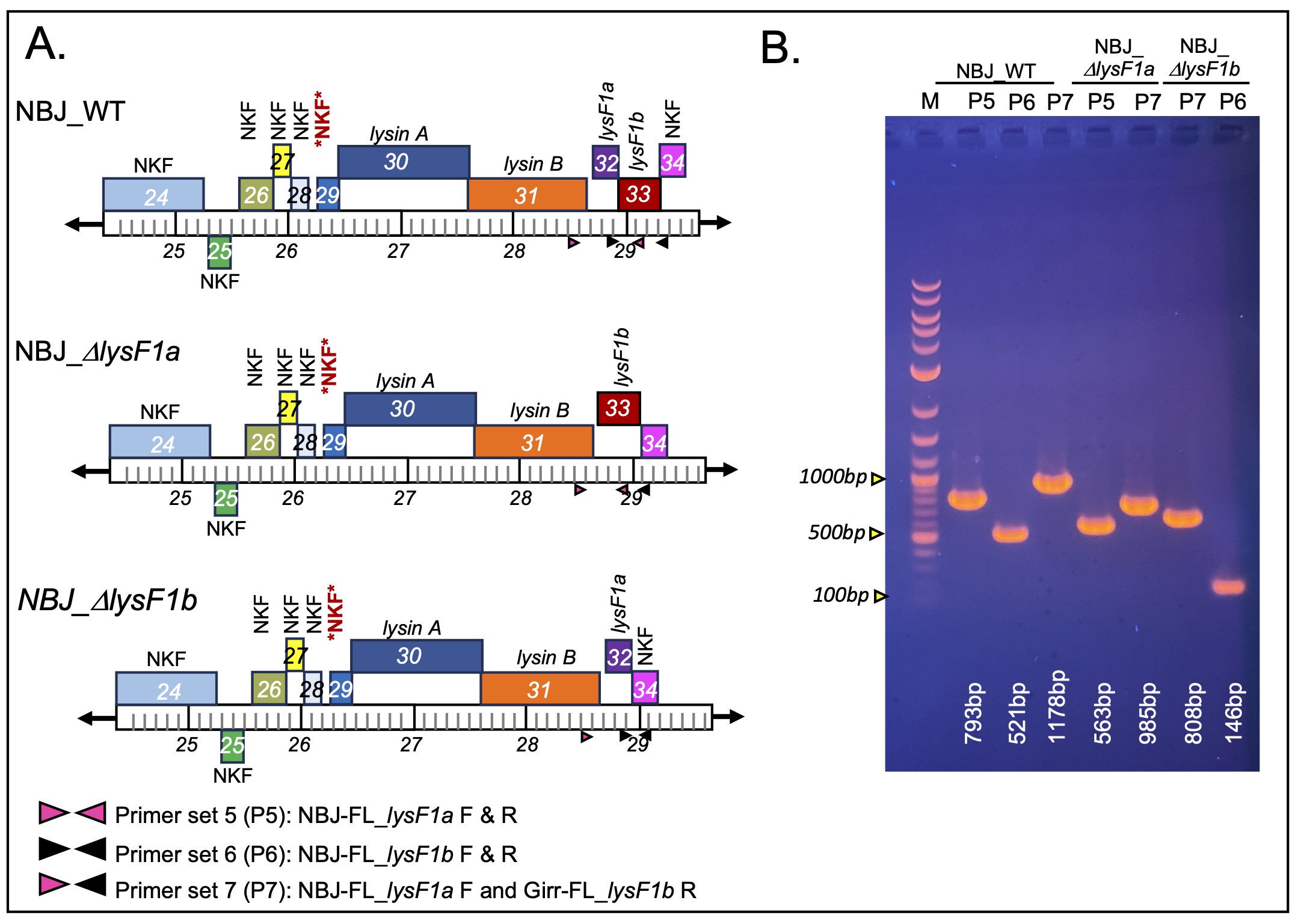

### S8 Figure 7

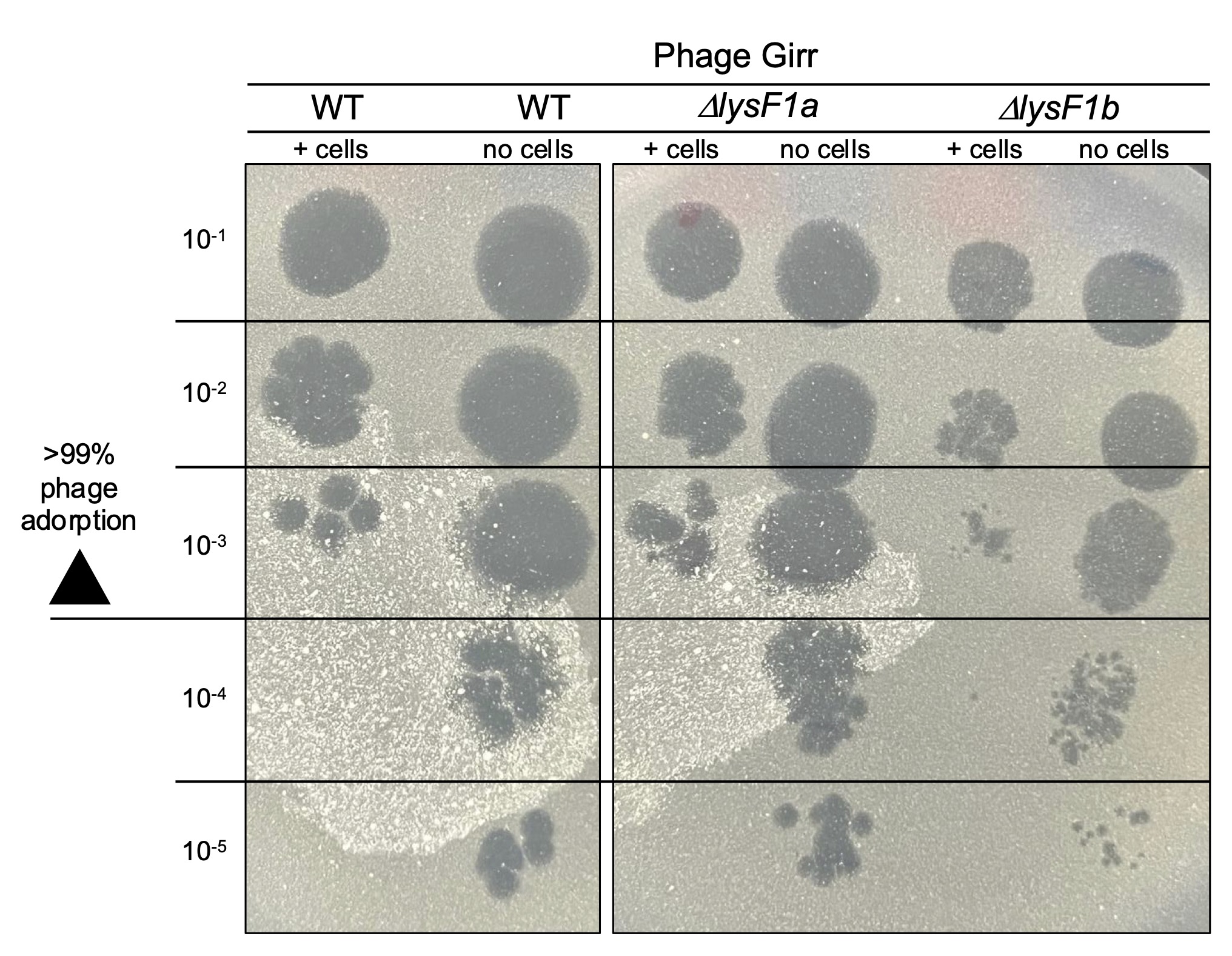

### S9 Figure 8

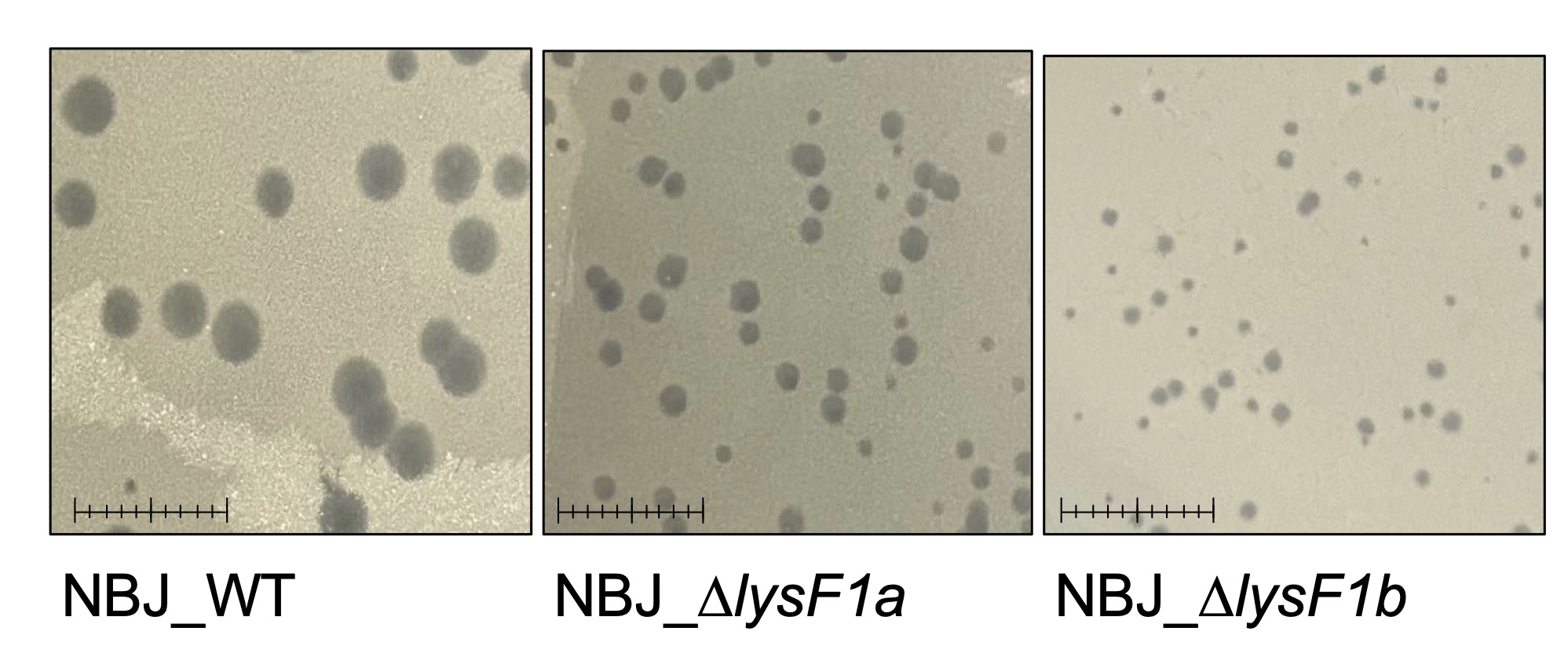

### S10 Figure 9

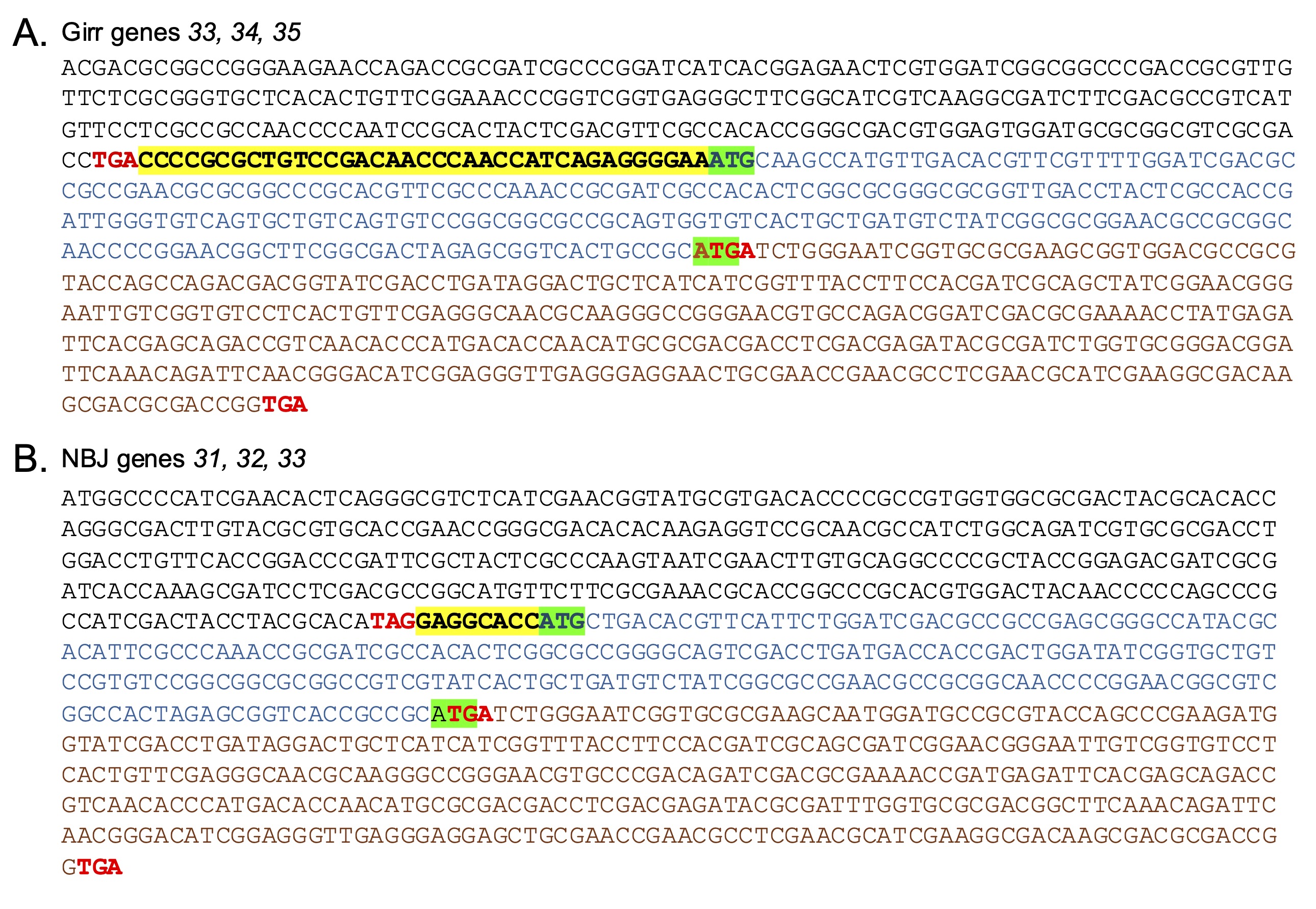

### S11 Figure 10

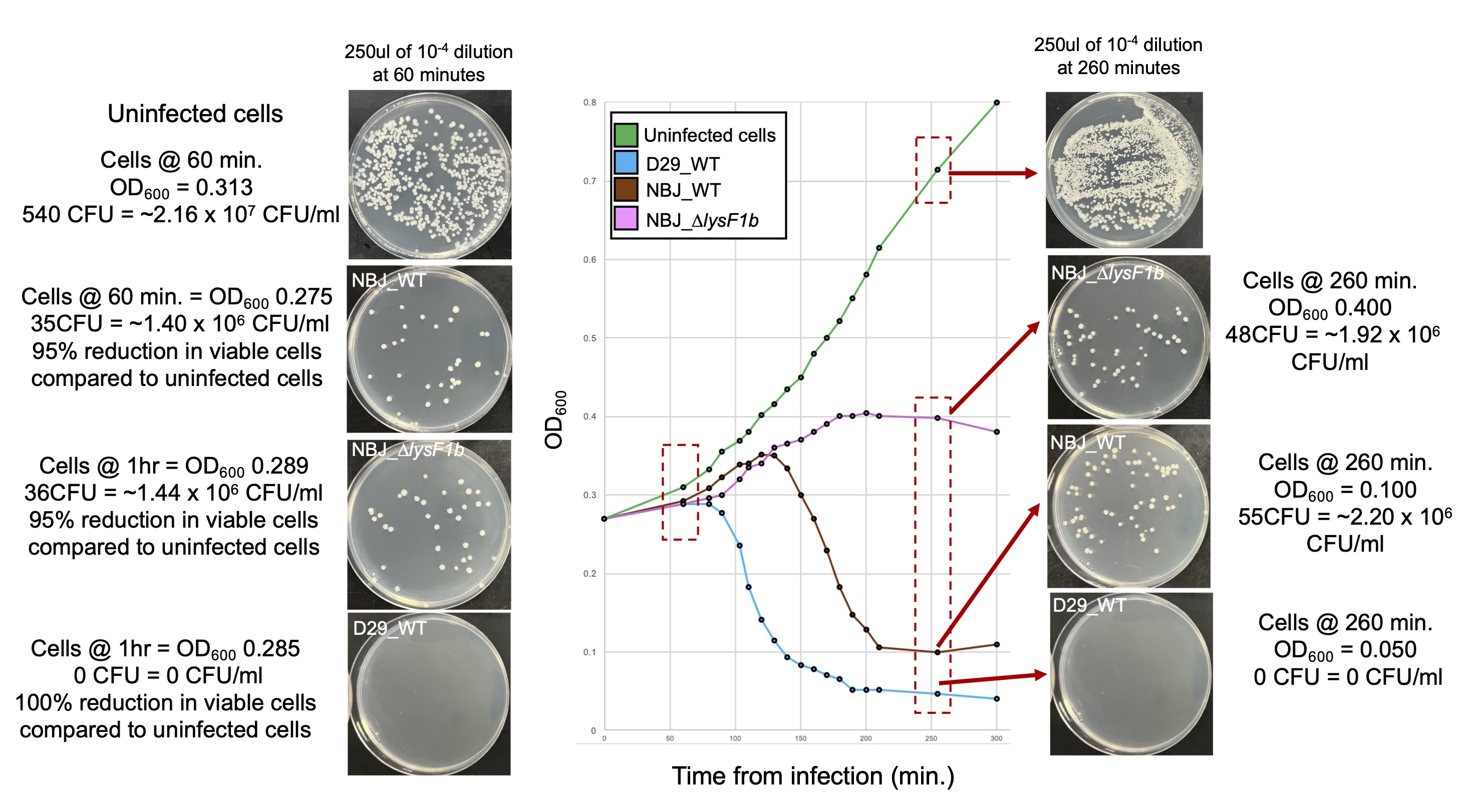

### S12 Figure 11

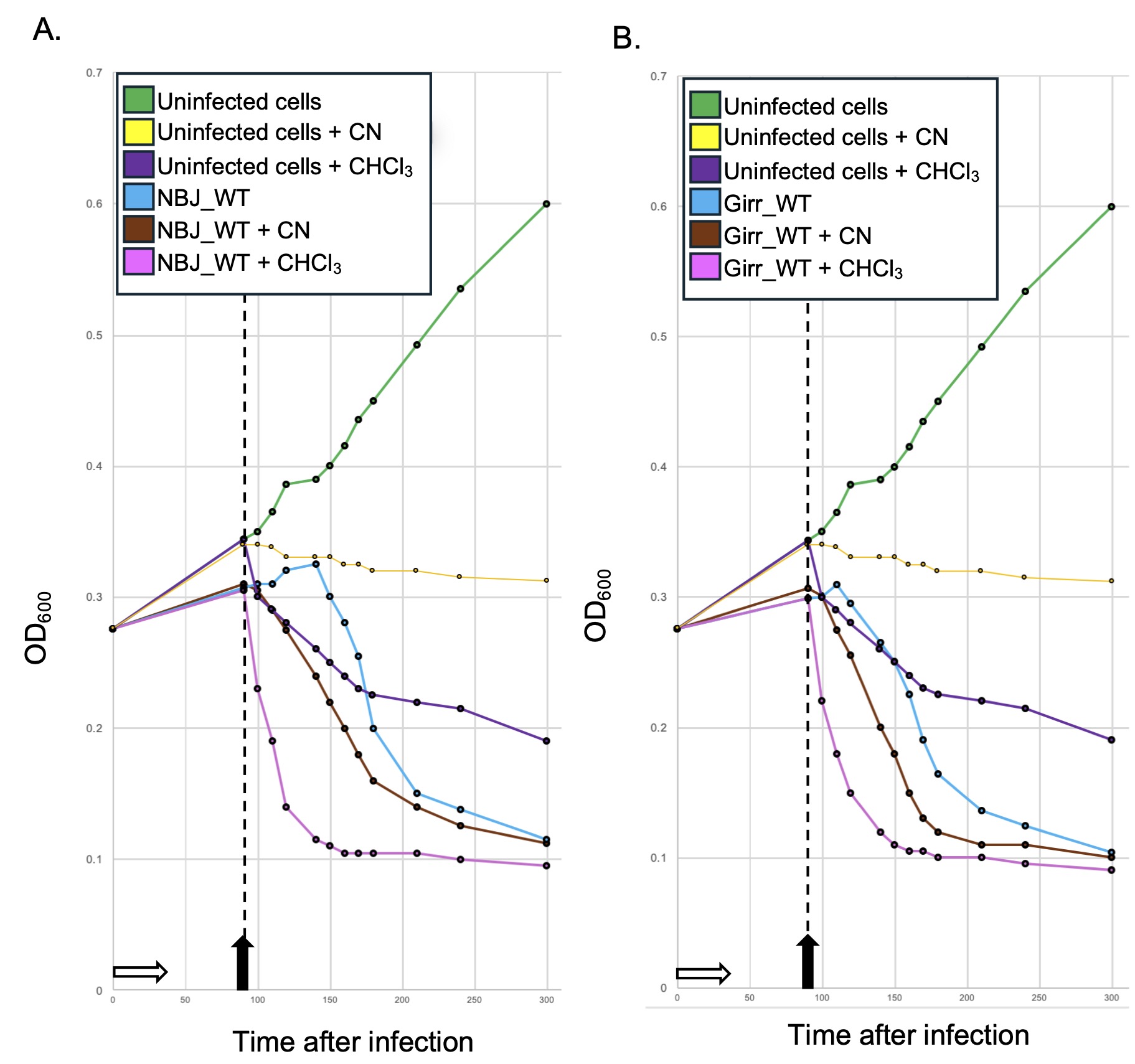

### S13 Figure 12

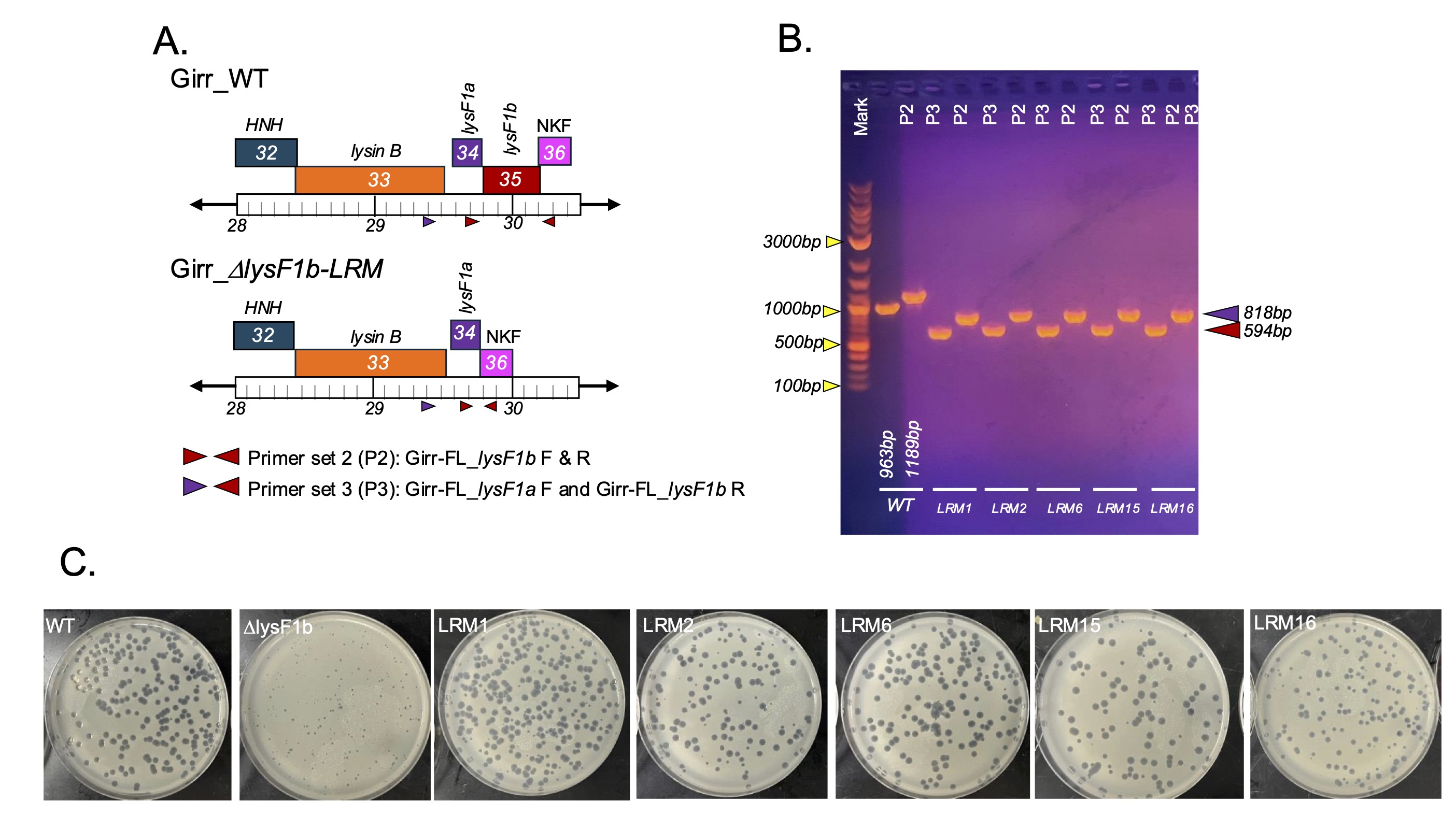

### S14 Figure 13

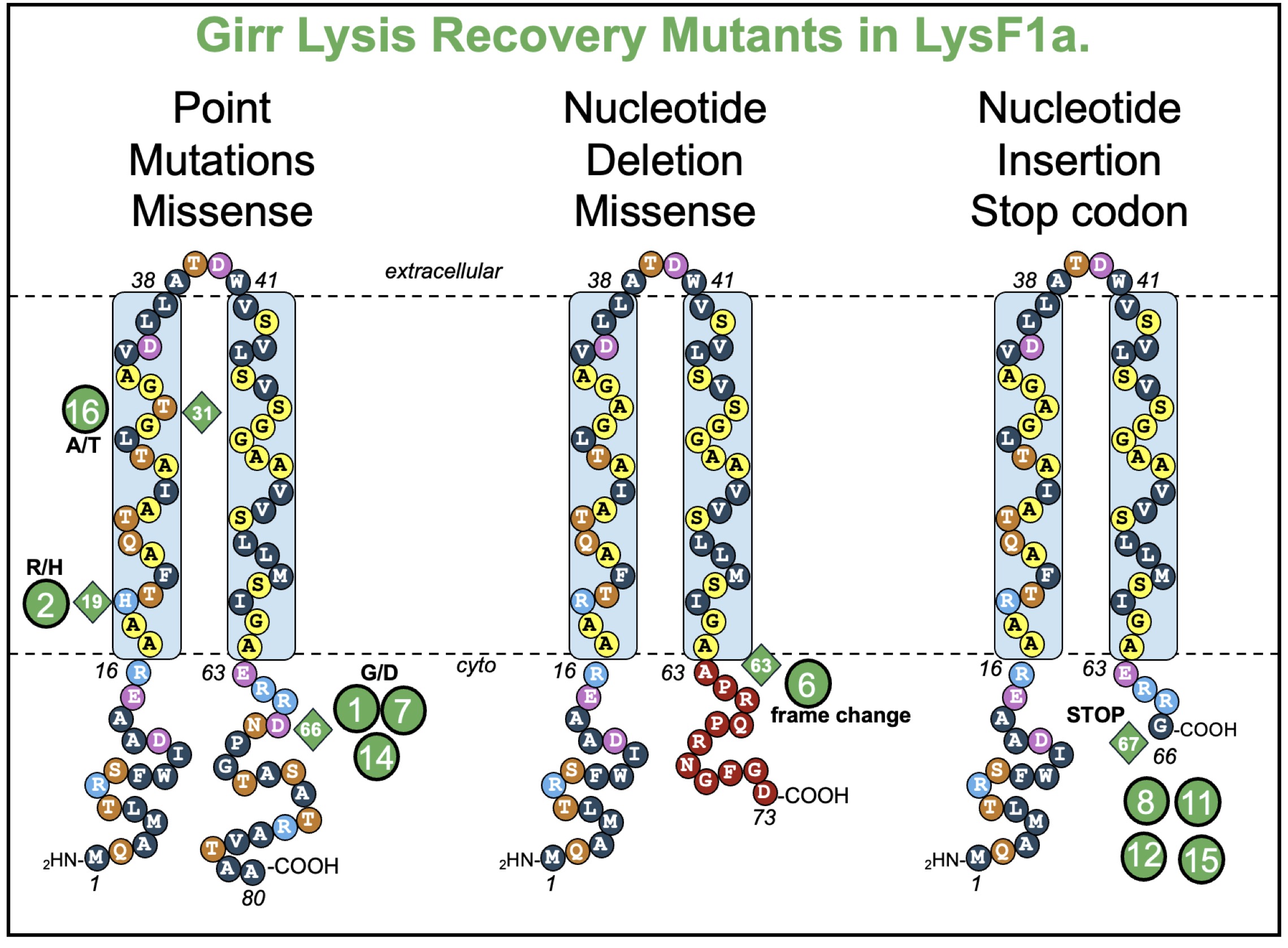

### S15 Figure 14

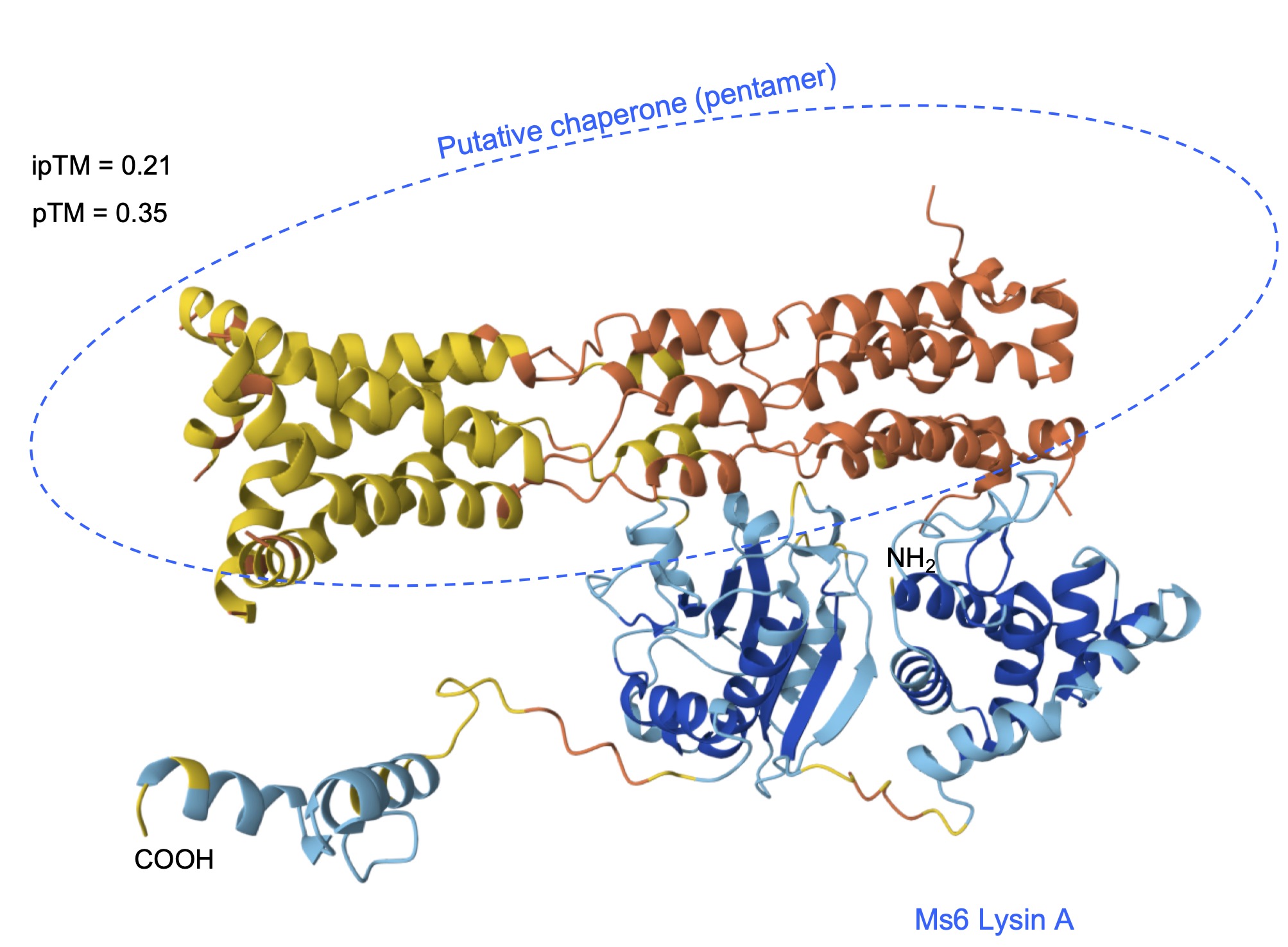

### S16 Figure 15

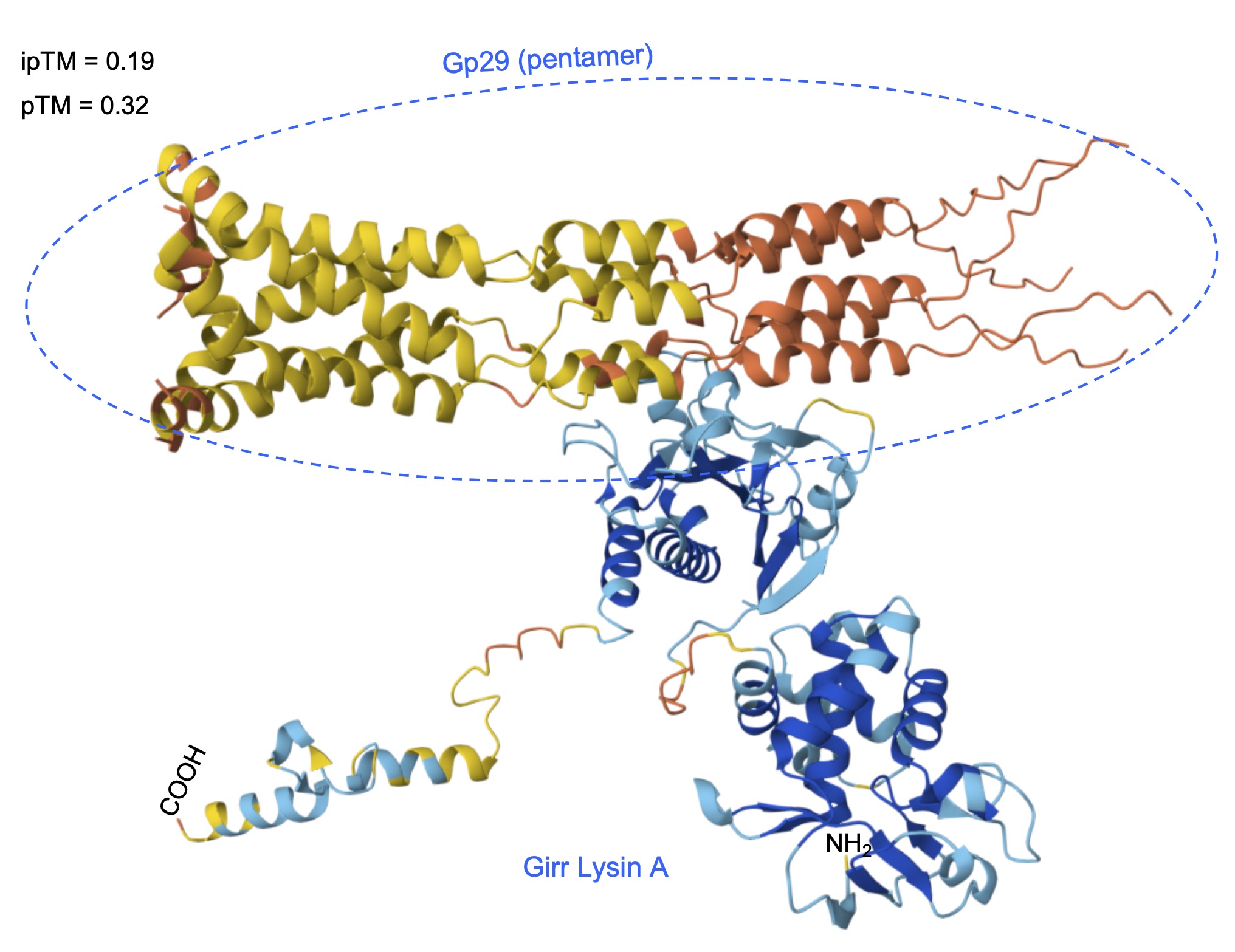

### S17 Figure 16

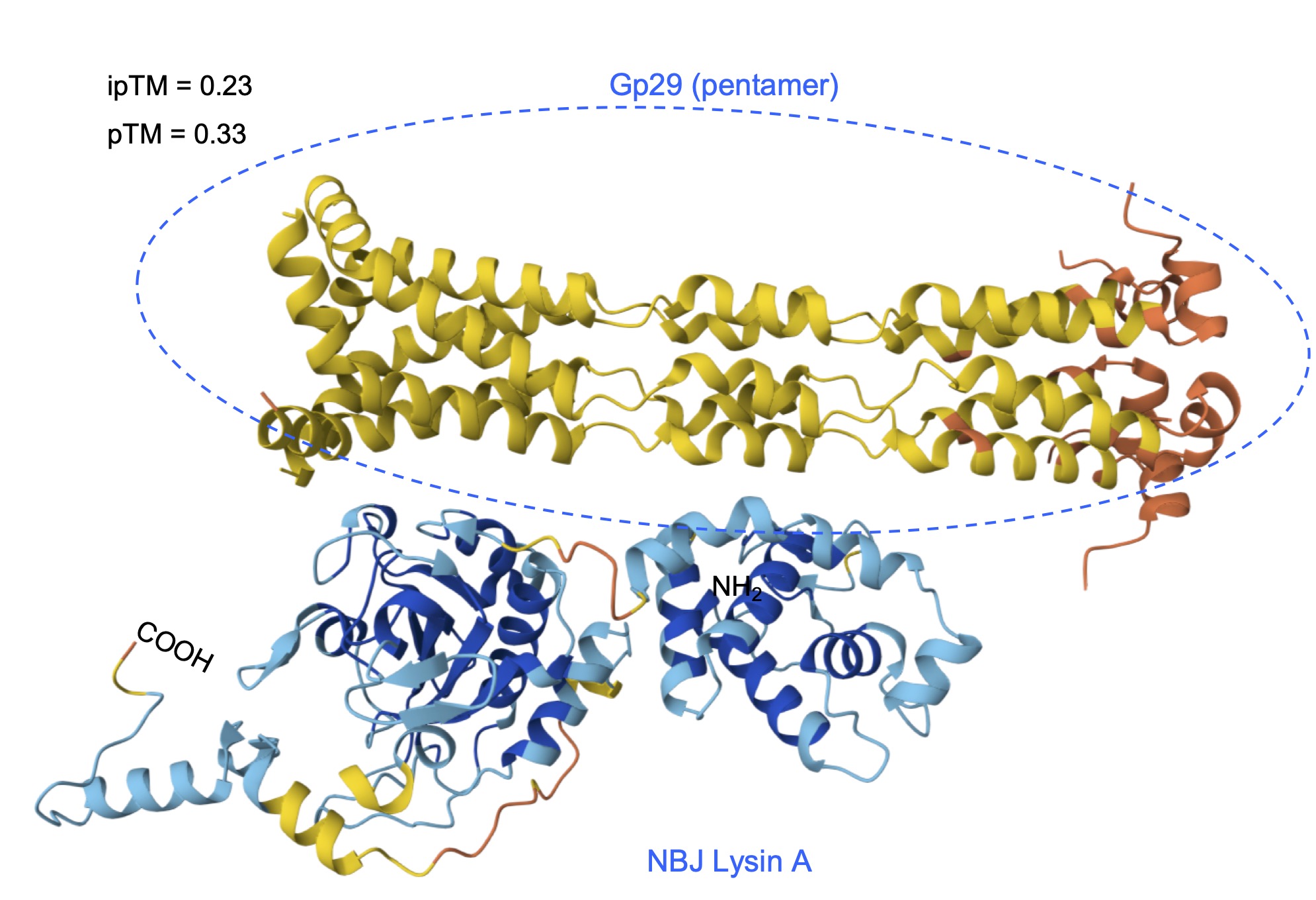

### S18 Figure 17

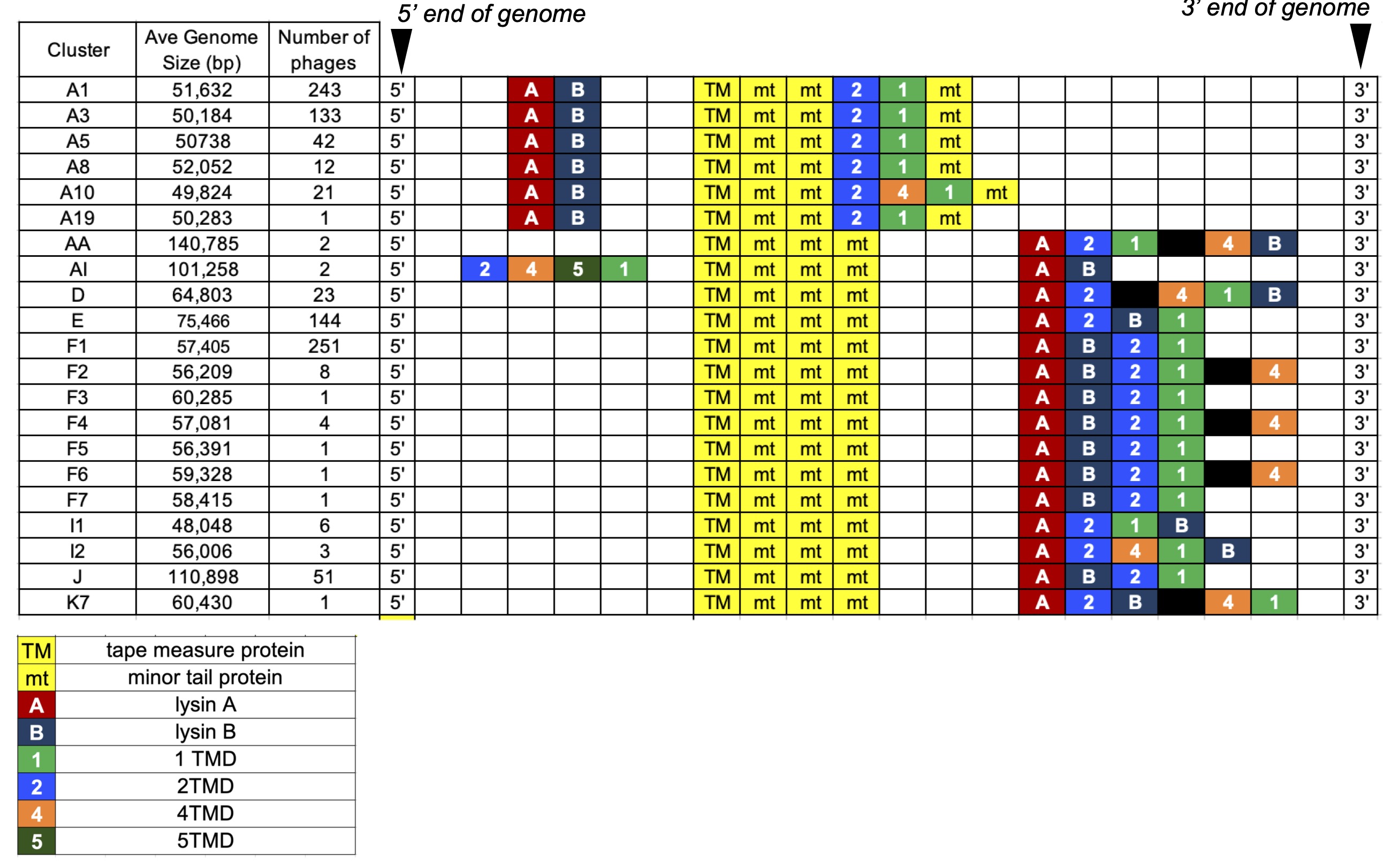

### S19 Figure 18

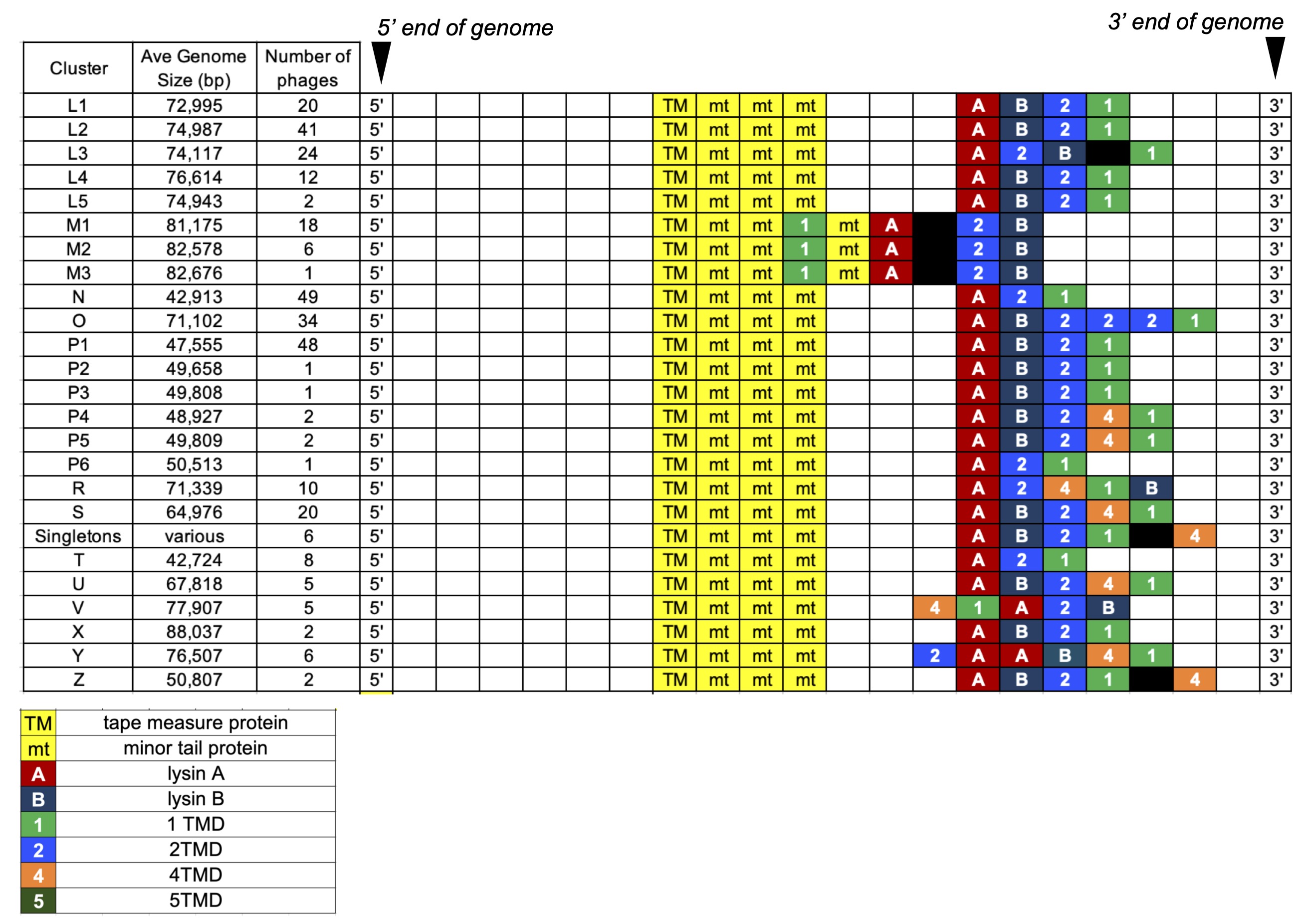

### S20 Figure 19

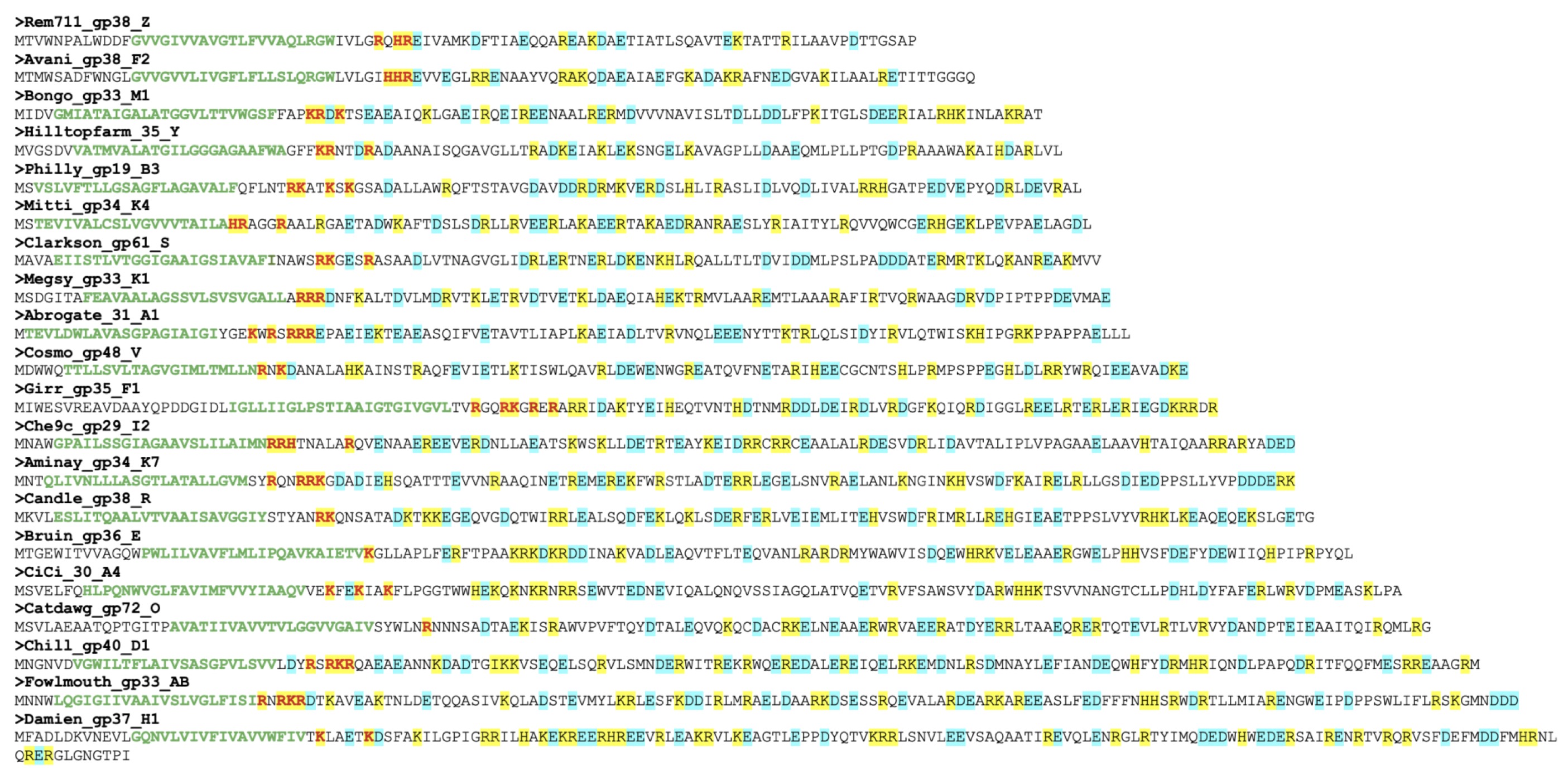

### S21 Figure 20

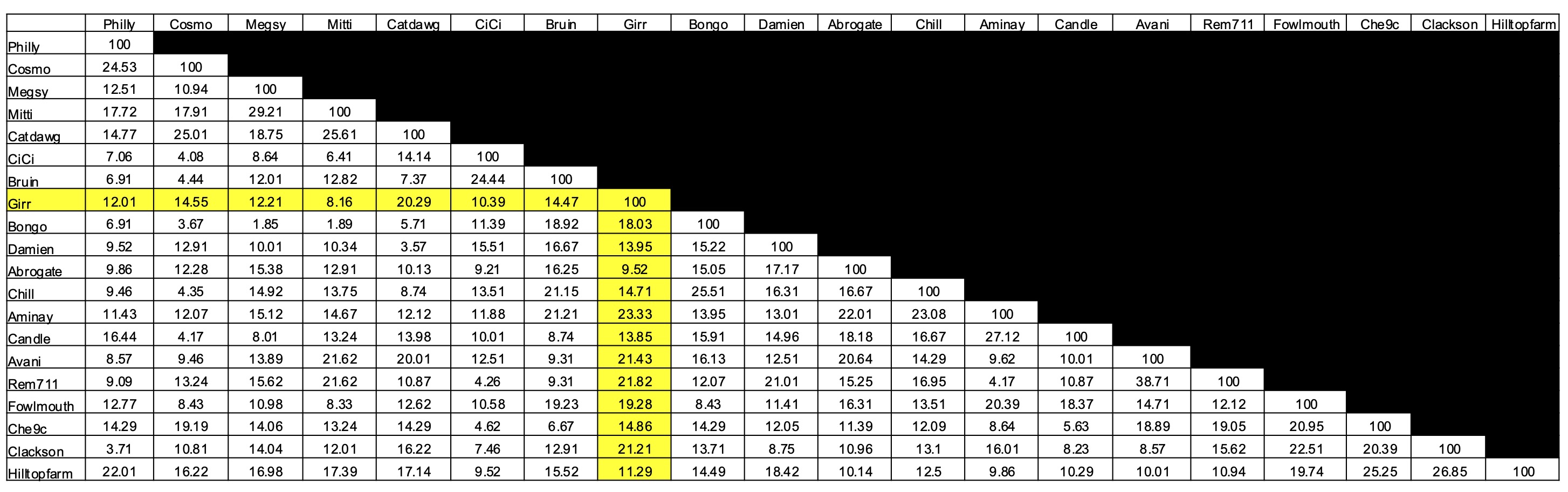
